# Resomapper: A Versatile Pipeline for Multiparametric MRI Processing. A Demonstrative Application in Mouse Brain Imaging

**DOI:** 10.1101/2025.08.20.671222

**Authors:** Raquel González-Alday, Adriana Ferreiro, Nuria Arias-Ramos, Blanca Lizarbe, Pilar López-Larrubia

## Abstract

**Objectives:** Magnetic resonance imaging (MRI) is essential in both research and clinical settings, with quantitative MRI (qMRI) enhancing reproducibility and sensitivity. However, qMRI processing can be complex, especially for users with limited coding experience. We introduce Resomapper, an open-source, cross-platform tool that integrates established processing libraries into a unified, user-friendly workflow, simplifying qMRI analysis while promoting accessibility, reproducibility, and data sharing.

**Materials and Methods:** Resomapper is a Python-based pipeline designed for intuitive multiparametric MRI processing. It supports T_1_, T_2_, and T_2_* relaxometry, magnetization transfer imaging (MTI) and diffusion tensor imaging (DTI) model fitting. The software includes advanced preprocessing options such as denoising, Gibbs artifact removal, and bias field correction. Users can process data interactively through a sequential pipeline or via an automated JSON-configured workflow, both executable through simple command-line instructions. Resomapper ensures compatibility by converting raw MRI data from different formats into the standardized NIfTI format within a BIDS-like structure, enhancing reproducibility, scalability, and data management efficiency. To demonstrate its application, we present a brain MRI study carried out on healthy, adult C57BL/6J mice, both sexes. MRI acquisitions were conducted on a Bruker Biospec 7T system using a multiparametric MRI protocol that included anatomical T_2_W images, T_2_ and T_2_* maps, MTI and DTI. The data were processed with Resomapper and co-registered using ANTsPy. Finally, a region of interest (ROI)-based analysis was performed to examine differences between sexes and brain areas, focusing on the cortex (Cx), hippocampus, (HPC), thalamus (Thal), and hypothalamus (HTH).

**Results:** Differences in all MRI parameters were found across brain regions, as expected. Additionally, a small significant sex difference in T_2_* was observed, with higher values in the thalamus and hypothalamus of female mice compared to males. This may reflect sex-specific responses to anesthesia. Moreover, this study also serves as a reference for standardized multiparametric qMRI studies in mice using Resomapper.

**Conclusions:** By integrating multiple processing tools into a single, accessible framework, Resomapper streamlines qMRI workflows and enables reproducible, high-quality image analysis. Thanks to its ability to handle diverse preprocessing techniques, multiple imaging modalities, and standardized data formats, the software proves to be a valuable resource for researchers with varying levels of programming expertise.

## 1. Introduction

Magnetic resonance imaging (MRI) is a highly valuable tool for both clinical diagnosis and research, offering detailed, non-invasive insights into tissue structure and function. In clinical practice, MRI is typically interpreted qualitatively, with radiologists usually assessing contrast variations across differently weighted images. However, these images are influenced not only by the underlying biophysical phenomena they are intended to highlight, but also by extrinsic factors, such as scanner-specific settings and hardware differences. These confounding influences limit the suitability of conventional MRI for quantitative analysis and hinder reproducibility across subjects, time points and imaging centers (1,2). To overcome these limitations, quantitative MRI (qMRI) techniques have been developed to extract direct, biophysically meaningful measurements of tissue properties. By minimizing the impact of extrinsic factors, qMRI improves reproducibility and reduces bias, thereby enhancing the ability to detect subtle microstructural or functional abnormalities that might remain undetected with qualitative approaches (1). Thanks to its increased sensitivity and interpretability, qMRI has become an indispensable tool in both clinical and research applications. Nevertheless, challenges remain, particularly the need to optimize acquisition times and to ensure consistent experimental conditions, which are critical to reducing variability in parameter estimation. Despite these challenges, the non-invasive nature, high precision, and three-dimensional spatial resolution of qMRI underscore its potential as a powerful tool in research and translational medicine (3,4).

A wide range of qMRI techniques are available, including relaxometry (quantification of T_1_, T_2_ and T_2_* relaxation times) (4); magnetization transfer imaging (MTI) (5), chemical exchange saturation transfer (CEST) imaging (6); diffusion weighted imaging (DWI), encompassing techniques like diffusion tensor imaging (DTI) (7) and diffusion kurtosis imaging (DKI) (8); functional MRI (fMRI) (9); quantitative susceptibility mapping (QSM) (10); magnetic resonance spectroscopic imaging (MRSI) (11); and perfusion imaging techniques, such as arterial spin labelling (ASL) (12), dynamic susceptibility contrast (DSC) (13) or dynamic contrast-enhancement (DCE) (14). These methods typically involve specialized acquisition protocols in which acquisition variables are systematically modulated to model specific biophysical properties. Under physiological and pathological conditions, the parameters obtained via qMRI may serve as biomarkers, enhancing diagnostic sensitivity, supporting prognostic assessment, and enabling preclinical or clinical therapy monitoring.

Following acquisition, generating the final parametric maps requires multi-step processing pipelines, which involve specific model fitting procedures and often extensive pre- and post-processing algorithms for noise reduction, artifact correction, and image registration. This process can be particularly challenging for researchers with limited programming experience. Moreover, the diversity of available tools, often fragmented, modality-specific, or incompatible with certain operating systems or data formats, further complicates software selection. The frequent reliance on non-public in-house software and lack of standardization also hinders reproducibility and transparency in qMRI research. Taken together, these factors highlight the urgent need for user-friendly, open-source processing tools that simplify complex workflows and enable a wider research community to take full advantage of qMRI. Although many excellent open-source tools have been developed for qMRI data processing—including dipy (15), DSI Studio (16), MRtrix (17), FSL (18), SPM (19), and AIDA MRI (20)—most of them are tailored for specific modalities, particularly DWI or fMRI. While more comprehensive frameworks such as qMRLab (21), support multiple modalities, they often require complex installations or rely on proprietary platforms like MATLAB, introducing additional barriers. To overcome these limitations, we introduce Resomapper, a cross-platform, open-source, pipeline that supports multiple qMRI modalities and simplifies data conversion and processing within a BIDS-like structure using the NIfTI format. Resomapper integrates several established open-source tools into a streamline, organized, and user-friendly Python-based workflow. It requires no compilation, it is easy to install, and it is adaptable to different research needs. Its accessible code also allows users with limited programming expertise to understand, apply and extend its functionalities. By promoting standardized data structures, enhancing reproducibility, and simplifying complex workflows, Resomapper provides an accessible and versatile solution for the research community. This paper presents an overview of Resomapper and demonstrates its application in a multiparametric MRI study of the mouse brain, focusing on sex-based and regional differences in C57BL/6J mice. While the case study involves preclinical data, Resomapper is equally is equally applicable to clinical datasets or preclinical data from other species, highlighting its versatility and broad applicability in qMRI research.

## 2. Software Overview

### 2.1 General design principles

Resomapper is a Python-based software package designed to streamline the analysis of multiparametric MRI data. The source code is freely accessible on GitHub (https://github.com/Biomedical-MR/resomapper) and can also be installed via the Python Package Index (PyPI), allowing users to either employ the distributed version or modify the source code to meet specific research needs. Python, known for its code readability and user-friendly nature, is particularly well-suited for researchers with limited coding experience. As an interpreted language, it eliminates the need for compilation, thereby simplifying installation, usage, and sharing of scripts and packages. This accessibility makes Resomapper straightforward to use, highly adaptable, and compatible across various operating systems. The software can be installed effortlessly using the package installer for Python (Pip), and detailed documentation is hosted online (https://resomapper.readthedocs.io/en/stable/). The main objective of Resomapper is to integrate well-established libraries into a unified toolbox, providing a standardized and user-friendly interface for analysing multiparametric MRI data. This approach simplifies the processing workflow while maintaining robust functionality. The package relies on several prominent Python libraries to achieve its objectives. Specifically: Nibabel (22) handles reading, writing, and manipulating MRI data in NIfTI format. SimpleITK (23) facilitates resampling and a variety of image processing tasks. Dipy (15) supports DTI model fitting, denoising, and related functionalities. Relaxometry map fitting is adapted from the myrelax package (24). Matplotlib (25) is used for graphical representations, while NumPy (26) and SciPy (27) are integral to mathematical computations and model fitting. Additionally, the BrkRaw (28) and pydicom (29) packages enable the conversion of Bruker raw and DICOM data, respectively, to the NIfTI format. Currently, Resomapper supports the processing of T_1_, T_2_, and T_2_* relaxometry, as well as MTI, DTI, and classical apparent diffusion coefficient (ADC) fitting. The software is actively maintained, with plans to incorporate additional functionalities in future releases. It is designed for beginners and offers a step-by-step interface that allows users to closely supervise the processing workflow. Its intuitive design ensures ease of use, making it suitable for both exploratory or fine-tuned analysis.

### 2.2 Processing workflow

Resomapper offers two flexible processing options to accommodate users with varying levels of expertise and diverse analytical needs. As a Python package, its functions can be accessed through its API, enabling users to develop custom scripts tailored to their specific requirements. In addition, Resomapper includes two pre-implemented workflows: 1) an interactive sequential pipeline that is intuitive to use for beginners enabling users to closely supervise the processing workflow 2) an automatic workflow where users specify all necessary parameters in advance via a JSON configuration file. This approach is particularly suited for batch work and parallel computing. Both workflows can be executed via simple command-line instructions, providing accessibility to users across different technical backgrounds. Regardless of the chosen approach, the pipeline adheres to a standardized sequence of steps, shown in Fig 1 and detailed in the subsequent sections.

**Fig 1.**
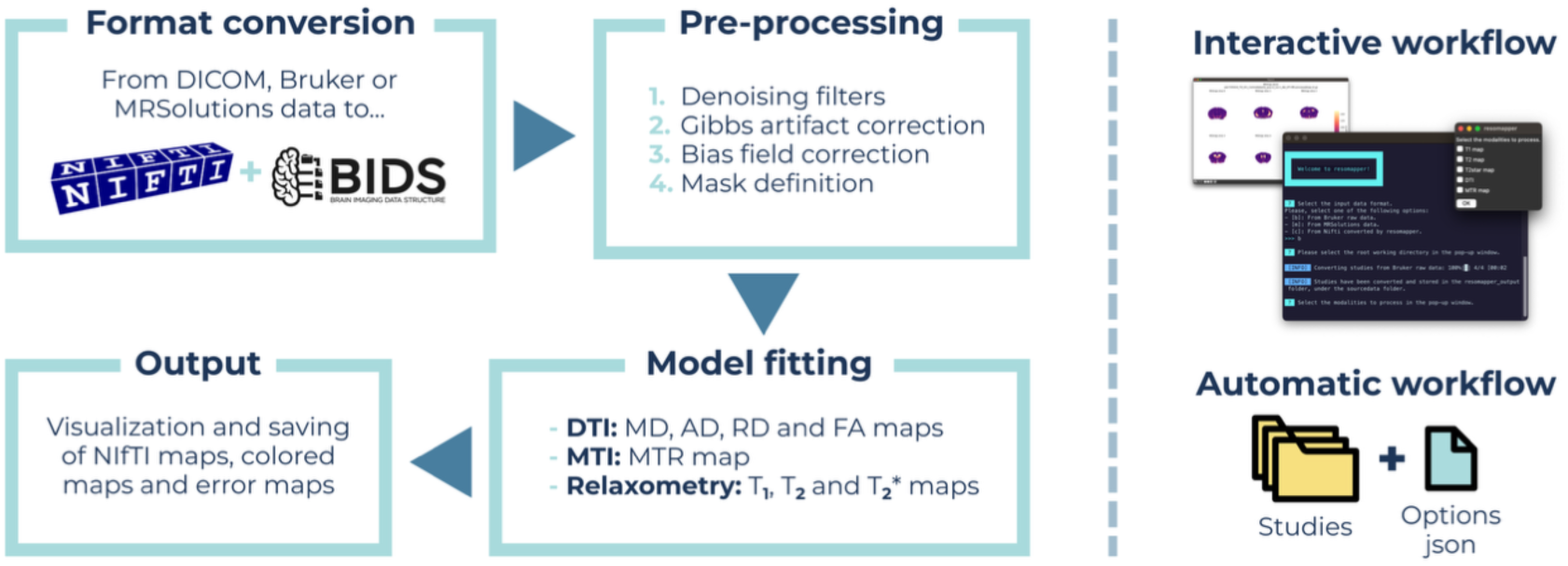
Overview of the steps implemented in the processing pipeline of Resomapper. This process can be accessed from either an interactive workflow, controlled sequentially via questions and graphical windows, or from an automatic workflow.

#### 2.2.1 Format conversion

MRI data can be stored in various vendor-specific raw formats, creating challenges for standardization and data handling. Resomapper has been designed to utilize the NIfTI (30) format due to its open-source nature, ease of handling, compact storage requirements, and capability to store of 3D or 4D image stacks. Metadata associated with each image, like the information typically found in DICOM headers, is stored in separated JSON files, which are easily readable text files structured as dictionaries facilitating the organization and retrieval of information. All image and metadata files are organized according to the Brain Imaging Data Structure (BIDS) (31), a widely adopted standard developed to promote consistent, intuitive organization and sharing of imaging data (Fig 2). The first step in the Resomapper workflow involves converting input data into this standardized format when necessary. The software currently supports conversion from three specific formats: DICOM data (32); raw data from Bruker preclinical MRI scanners (Bruker Medical GmbH^®^, Ettlingen, Germany); and data from MR Solutions preclinical scanners (MR Solutions Ltd., Guildford, UK). By converting data into the NIfTI format and adopting the BIDS organizational structure, Resomapper ensures compatibility, scalability, and efficient data handling for both preclinical and clinical applications.

**Fig 2.**
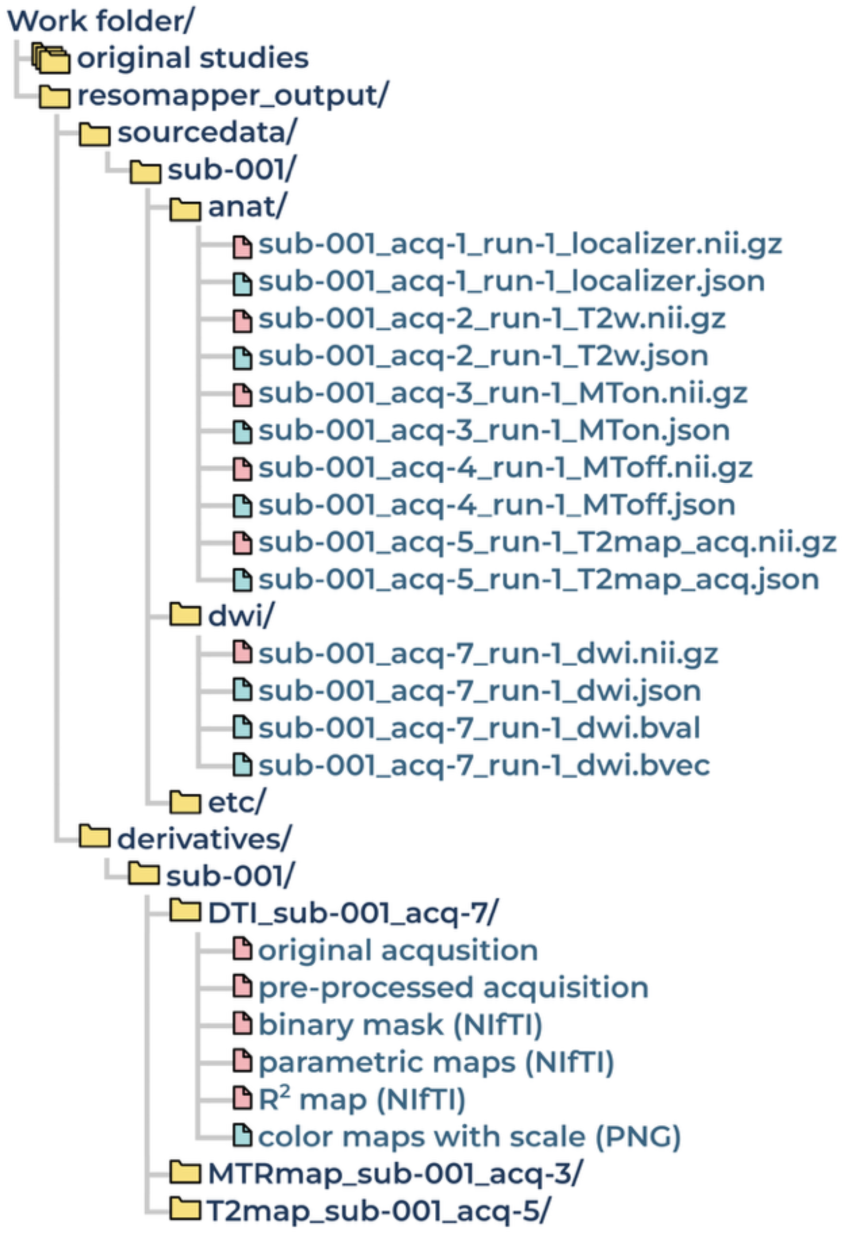
**Scheme of the BIDS-like structure designed for Resomapper output Eiles.**

#### 2.2.2 Pre-processing options

Before fitting the MRI data to the corresponding models and generating parametric maps, Resomapper offers a range of pre-processing options aimed at improving image quality. These tools are designed to reduce noise and correct certain artifacts, thereby enabling more accurate model fitting that results in reliable and precise parametric maps.

##### - Denoising filters

MRI data are susceptible to various sources of noise that can adversely affect the signal-to-noise ratio (SNR). These sources include body motion, physiological conditions, susceptibility effects at tissue interfaces, eddy currents, stochastic signal variations, and radiofrequency interferences (33). To address these issues, Resomapper incorporates several widely used denoising methods (Fig 3). These filters are particularly effective in reducing random noise, and include: non-local means filtering, from the scikit-image package (34); adaptive soft coefficient matching (ASCM); local principal component analysis (local-PCA); Marchenko-Pastur PCA (MPPCA), and Patch2Self, implemented through the Dipy library.

**Fig 3.**
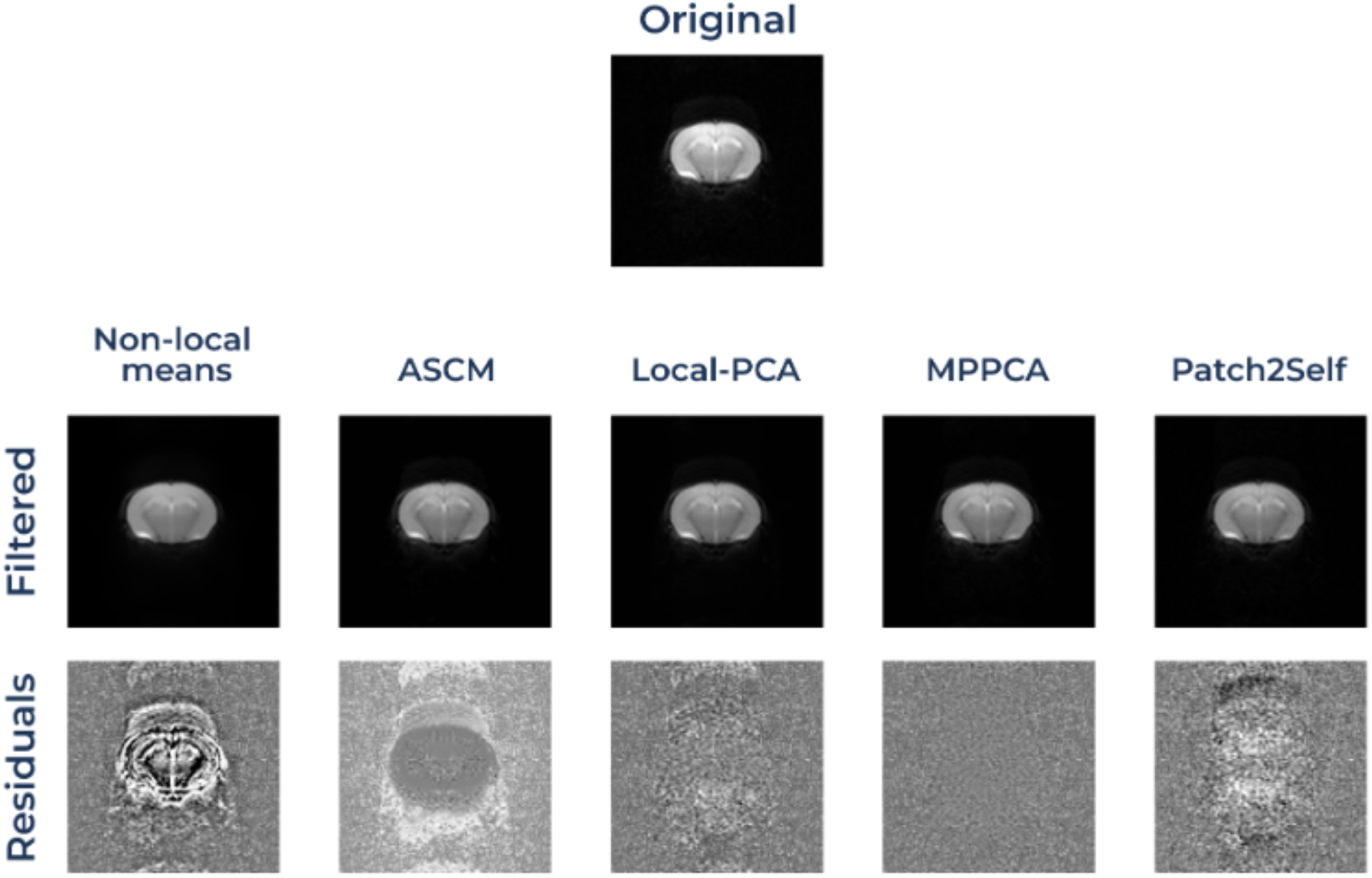
Example of the different denoising Eilters available in Resomapper. Filters were applied to a B_0_ image from a diffusion weighted acquisition. The filtered image and the residuals are shown to illustrate where has the image been altered. Note that, ideally, only random noise should be eliminated, but some filters like non-local means perform excessive smoothing, eliminating some details from the image.

The non-local means algorithm (35) reduces noise by replacing each pixel’s intensity with a weighted average of intensities from surrounding pixels. This process involves comparing small patches centered on surrounding pixels with the patch centered on the target pixel, so that pixels with patches that closely match the target patch are included in the averaging process.

ASCM (36) method is an extension of non-local means denoising that enhances image quality by combining two denoised versions of the original data with different levels of smoothness. Using wavelet decomposition, it merges low-frequency components (smoothness) from the more smoothed input with the high-frequency details (sharpness) from the less smoothed input, resulting in a well-balanced and enhanced denoised output.

PCA-based methods suppress noise by identifying and removing noise components from the data. These methods are particularly effective for reducing noise in diffusion-weighted or relaxometry data (37), as they leverage the inherent redundancy in their multi-dimensional structure. Principal component classification can be guided by prior noise variance estimates, as in the local-PCA filter (38), which estimates noise variance locally at each voxel, applies PCA to patches across gradient directions, and thresholds eigenvalues before reconstructing the data. Alternatively, the MPPCA (39) approach bypasses the need for prior noise variance estimation to automatically determine noise thresholds based on the Marchenko-Pastur distribution.

Finally, Patch2Self (40) is a self-supervised denoising algorithm specifically designed for DWI data. It leverages the entire dataset to construct a locally linear, full-rank denoiser uniquely adapted to that volume. By exploiting the dense sampling in DWI data, Patch2Self efficiently separates noise from structural information without requiring predefined models.

##### - Gibbs artifacts suppression

In addition to noise, a common artifact encountered in MRI are Gibbs artifacts, also known as ringing artifact. These arise during the reconstruction of the MR signal from the k-space using Fourier transforms. Briefly, the finite sampling of frequencies in the MRI k-space results in the truncation of the Fourier series, leading to an imperfect representation of the data. (41). Gibbs artifacts are particularly notable at interfaces with sharp intensity transitions, such as tissue boundaries, where this truncation leads to ring-like oscillations in the signal near the edges (42). Several approaches exist to mitigate these artifacts, including increasing spatial resolution, employing extrapolation methods like zero filling during reconstruction, or using smoothing filters. In Resomapper, Gibbs artifact suppression is addressed through Dipy’s implementation of a sub-voxel shift method. This approach reduces ringing with minimal image smoothing by locally shifting the image grid to align points with positions where the oscillatory pattern cancels out (43,44). This method provides an effective balance between artifact suppression and preservation of spatial detail (Fig 4).

**Fig 4.**
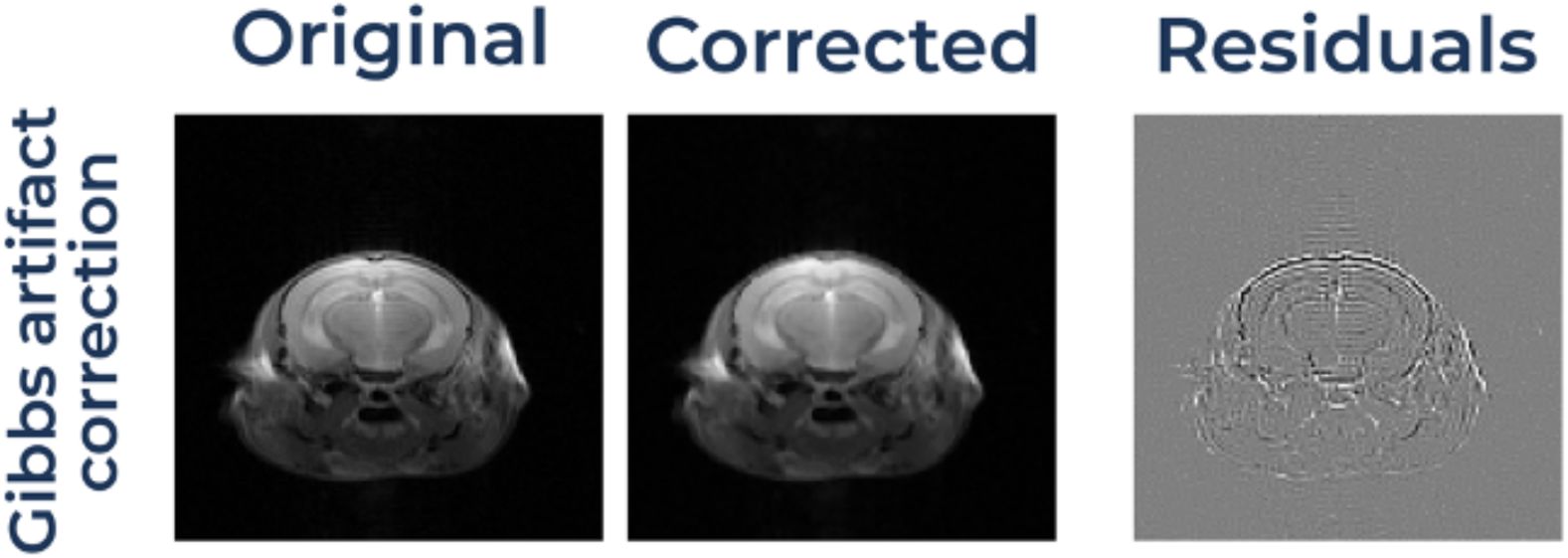
Example of the Gibbs artifact correction. Gibbs artifact correction applied to an MTon image, and the resulting residuals, where the ringing structures removed can be identified.

##### - Bias field correction

The bias field refers to low-frequency spatial variations in MRI image intensity that are unrelated to the underlying tissue characteristics. These variations can cause identical tissue types to appear with different intensity levels across the image, potentially affecting quantitative analyses. Bias field effects typically arise from non-uniformities in the magnetic field or from uneven sensitivity of radiofrequency coils, such as those in surface coils. A widely used method for correcting these intensity inhomogeneities is the N4 algorithm (45), a refinement of the earlier N3 (nonparametric nonuniform normalization) retrospective bias correction algorithm (46). It models the bias field using the image’s intensity histogram, iteratively adjusting intensity levels through a deconvolution operation, and smoothing the results spatially with a B-spline representation. Resomapper incorporates the SimpleITK implementation of the N4 algorithm to correct for bias field effects, thereby improving image uniformity and consistency in MRI image intensity (Fig 5).

**Fig 5.**
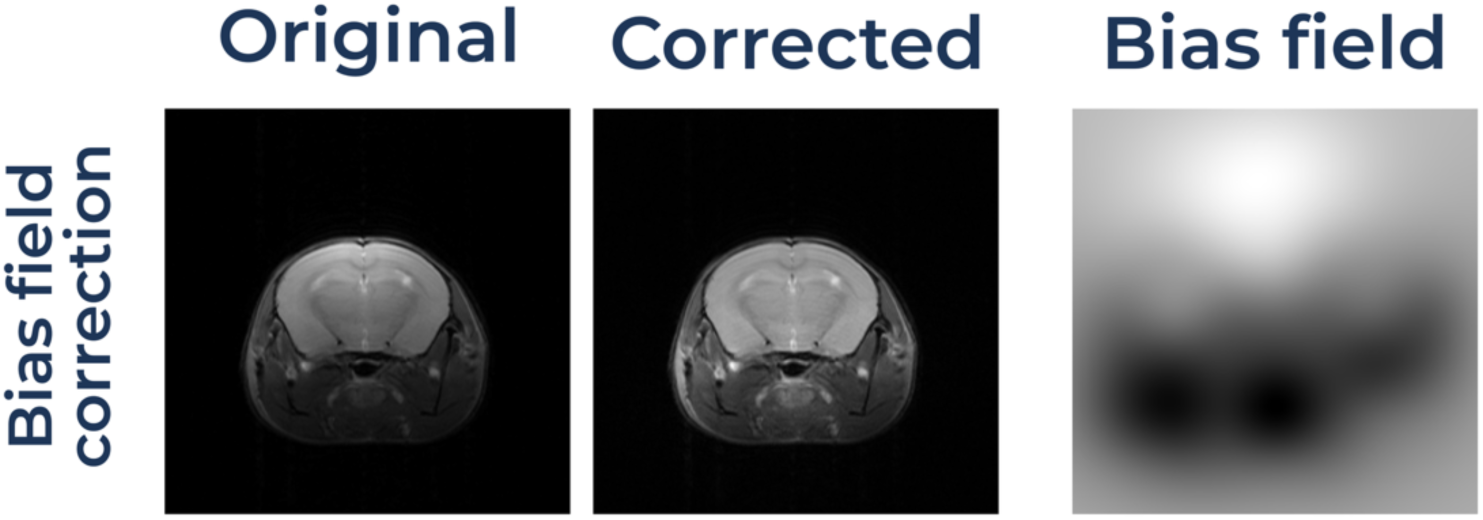
Example of the bias Eield correction applied to an anatomical T_2_W image. The panel shows the original T_2_W image, the corrected one and the bias field image.

##### - Mask definition

Defining a mask to delineate the region where the model fitting will be performed is particularly beneficial for relaxometry and DWI. This step reduces computational time, accelerating the processing workflow. Additionally, mask definition is advantageous when image co-registration, either between images or with an atlas, is intended. Resomapper currently offers four options for mask definition: i) users can manually draw the mask slice by slice, providing precise control over the selected region; ii) users can supply a binary mask file that specifies the region of interest; or iii) the masking step can be omitted entirely if not required for the analysis. These flexible options allow users to tailor the masking process to their specific needs, enhancing both efficiency and usability.

#### 2.2.3 Signal model fitting

After pre-processing the acquired images, parametric maps can be generated for further analysis. The following describes the processes and sequences required for each modality currently supported by Resomapper.

##### - Relaxometry

MRI relaxometry encompasses techniques used to quantitatively measure tissue relaxation properties, specifically the relaxation times T_1_, T_2_, and T_2_*. These parameters describe the behaviour of proton magnetization following exposure to an external magnetic field and the application of radiofrequency (RF) pulses. Since relaxation times are influenced by the biophysical and biochemical properties of tissues, they serve as valuable biomarkers for tissue characterization (4). Relaxometry maps, representing relaxation times on a pixel-by-pixel basis, are generated by fitting sets of images acquired at varying echo times (TE) or repetition times (TR) to the corresponding signal decay equations (47).

T_1_ maps measure the longitudinal relaxation time, which reflects how protons realign with the main magnetic field after excitation by an RF pulse. T_1_ is sensitive to molecular interactions and tissue composition, providing insights into different tissue types and pathologies, especially in tumors (48) and myocardial affections (49). T_1_ mapping commonly employs saturation-recovery sequences variable TR, where obtained signal intensities are fitted to the equation:

T1 mapping techniques often utilize inversion-recovery sequences due to their widespread adoption and established accuracy in clinical and research settings. However, in our software, we have implemented T1 mapping based on saturation-recovery sequences with variable TR. This approach offers notable benefits for animal studies, particularly in small animal models where physiological conditions such as rapid heart rates can pose challenges. In this case, signal intensities obtained at multiple TR values are fitted to the equation:

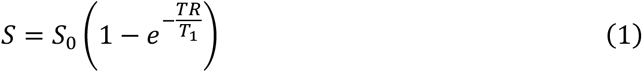

T_2_ relaxation quantifies the time required for protons to lose phase coherence in the transverse plane due to spin-spin interactions. This parameter is influenced by water content and tissue microstructure, making T_2_ measurement particularly useful for detecting pathologies like edema, inflammation or necrosis (50). T_2_ mapping typically uses multi-echo spin echo sequences, which acquire data at various TE with a fixed TR. The signal decay is modeled by fitting to the following equation:

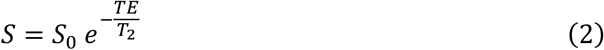

T_2_*, in contrast, measures effective transverse relaxation, encompassing both spin-spin interactions and magnetic field inhomogeneities. Unlike T_2_ mapping, which relies on spin echo sequences with a 180° refocusing pulse, T_2_* mapping typically uses gradient echo sequences, which lack this refocusing step. As a result, T_2_* is sensitive to additional dephasing caused by magnetic susceptibility differences or field distortions. This makes T_2_* mapping particularly well-suited for detecting magnetic or paramagnetic substances, such as iron, deoxygenated blood, or calcifications. It is therefore useful in studying conditions like hemorrhages or iron deposition (51–53). Signal intensities acquired at multiple TE using gradient echo sequences are fitted to the equation:

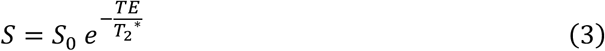

In Resomapper, the fitting of the multi-repetition times or multi-echo times data to the respective signal decay equations is accomplished using non-linear least squares optimization, with code adapted from the implementation described in (24).

##### - Magnetization transfer imaging

MTI is based on the exchange of magnetization between two distinct proton pools: the protons of free water molecules and those of water molecules bound to macromolecular structures with highly restricted motion (54). The bound proton pool cannot be directly detected in MRI due to its extremely short T_2_ relaxation time. However, these protons experience a heterogenous magnetic environment due to their restricted motion and interactions with macromolecules making them sensitive to a broader range of frequencies. This sensitivity enables selective excitation using an off-resonance RF pulse, which does not directly affect free water protons. When the bound proton pool is selectively excited, its magnetization transfers to nearby free water protons through dipolar and chemical exchange mechanisms. This process partially saturates the free proton pool and accelerates T_1_ relaxation, resulting in a reduced observable signal. To quantify this magnetization transfer effect, the magnetization transfer ratio (MTR) can be calculated using the formula:

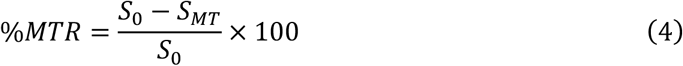

where S_MT_ is the observed signal when the magnetization transfer pulse is applied, and S_0_ is the observed signal in an equivalent image without this pulse.

MTR maps are sensitive to myelin content, and therefore very useful to assess lesions in neurodegenerative diseases like multiple sclerosis (55), as well as other abnormalities associated with edema, infection or tumors (56).

##### - Diffusion tensor imaging

DWI techniques rely on the diffusion of water molecules to generate contrast in MRI images. This is accomplished using fast T_2_-weighted (T_2_W) imaging through the application of two symmetric and apposite diffusion-sensitizing gradients before and after the 180-degree refocusing pulse. These gradients cause spin dephasing and subsequent rephasing, but rephasing is incomplete for protons that have moved during the interval between gradients. Complete rephasing only occurs for protons that remain stationary; protons that have moved significantly lose coherence, resulting in reduced signal intensity (57,58).

DWI sequences are based on the framework proposed by Stejskal and Tanner (59), with the associated signal attenuation described by the Stejskal-Tanner equation:

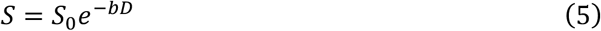

where S_0_ is the signal obtained in an identical acquisition without diffusion gradients, D is the apparent diffusion coefficient (as described by Einstein’s equation, assuming free diffusion) and b is a parameter that describes the diffusion weighting and is defined as:

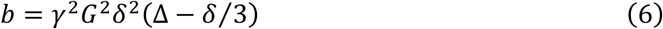

In this equation γ is the gyromagnetic ratio, G is the gradient amplitude, δ is the gradient duration and Δ is the time between the two gradients application.

By applying the Stejskal-Tanner equation and acquiring at least two diffusion-weighted images with two different b-values, the ADC can be estimated.

However, the Stejskal-Tanner assumes isotropic diffusion, representing unrestricted Brownian movement. In biological tissues, water movement is often hindered by microstructures, such as cell membranes, vessels and fibers. To better describe water diffusion in tissues advanced models have been developed, including DTI (60). DTI expands on DWI by modeling diffusion as a second order tensor, represented by a 3×3 symmetric matrix that describes the diffusion process at each voxel.

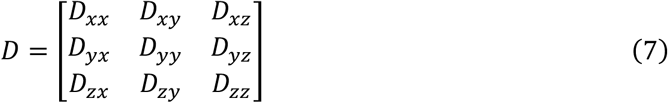

This tensor has six independent terms and can be visualized as an ellipsoid. Its estimation requires adapting the Stejskal-Tanner equation (see equation 8) and acquiring at least six diffusion-weighted images along non-collinear gradient directions, and at least one reference image without diffusion weighting (b = 0 s/mm^2^).

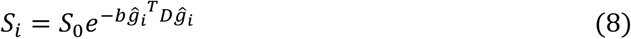

After the diffusion tensor is computed, it can be decomposed in three eigenvalues (ε_1_, ε_2_ and ε_3_) and their corresponding eigenvectors (λ_1_, λ_2_ and λ_3_), ordered by magnitude. From these, several diffusion metrics can be derived. These metrics are the mean diffusivity (MD), that measures the average diffusion along the three axes of the ellipsoid, axial diffusivity (AD), that reflects diffusion along the principal diffusion direction, radial diffusivity (RD), that quantifies diffusion perpendicular to the principal direction, and fractional anisotropy (FA) that describes the degree of diffusion anisotropy. FA is a scalar value ranging from 0 to 1. These metrics are calculated as:

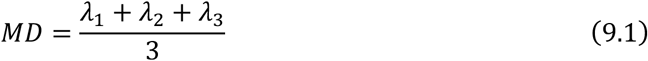

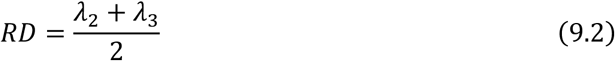

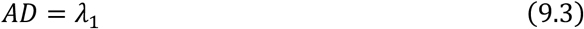

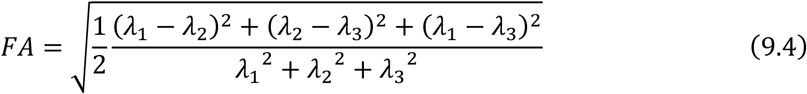

These metrics give insights about tissue microstructure, being commonly used to study neuronal damage, edema or neuroinflammation. Besides DTI metrics, Resomapper also supports the fitting of classical ADC from acquisitions with less than 6 gradient directions.

#### 2.2.4 Output saving

After processing, the parametric maps are stored in NIfTI format. In addition, the pre-processed original acquisitions and corresponding R^2^ (coefficient of determination) maps are saved to facilitate quality assessment and to check the goodness of fit. Furthermore, color-coded maps in PNG format are also generated for visualization purposes. The dynamic range and color scale of these images can be adjusted for each map in the final step of the workflow, according to user preference.

## 3. Materials and Methods

This section details the application of our software within a preclinical framework, where it was used to analyze multiparametric MRI data from the brains of male and female mice. By presenting this case study, we aim to exemplify the capabilities of the software in processing and extracting quantitative information from complex preclinical MRI datasets. Although no human MRI cases are included here, the described procedures and workflow are equally applicable and valuable for the analysis of human brain MRI data.

### 3.1 Animal models

All experimental procedures were performed in accordance with the national (R.D.53/2013) and European Community guidelines (2010/62/UE) for the care and management of experimental animals, and were approved by the Ethics Committee of the Community of Madrid (PROEX 288.4/22; approved 18 November 2021).

A cohort of 34 adult C57BL/6J mice, divided equally between males and females, was used in the study. Animals were housed in standard cages with ad libitum access to food and water at the Institute for the Biomedical Research Sols-Morreale (IIBM) animal facility (Reg. No. ES280790000188). Housed conditions included a 12 h light/dark cycle, a controlled temperature (22 ± 2 °C), and care by specialized personnel.

### 3.2 Magnetic Resonance Imaging

MRI acquisitions were carried out on a 7T scanner (BioSpec 70/16, Bruker Medical GmbH^®^, Ettlingen, Germany) equipped with a 23 mm receive-only surface coil and a 40 mm transmit-only resonator. The system incorporated a gradient insert of 90 mm in diameter and a maximum strength of 360 mT/m. All data were acquired running Paravision 6.0.1 software (Bruker Medical GmbH^®^, Ettlingen, Germany).

Mice were anesthetized with 3–4% isoflurane in 100% O_2_ in an induction chamber and subsequently positioned in an animal holder equipped with a heated blanket, to maintain body temperature at approximately 37 °C. A nose mask was used to deliver 1.5–2% isoflurane during MRI acquisitions. Vital parameters, including body temperature and respiratory rate, were continuously monitored and regulated using a monitoring and gating system (SA Instruments, Inc., Stony Brook, NY, USA).

The MRI protocol included T_2_W imaging and parametric MRI acquisitions (MTI, T_2_ mapping, T_2_* mapping and DTI). All images were acquired with a field of view (FOV) of 23×23 mm^2^, axial orientation, covering 5 slices of 1 mm thickness each, centred on the brain with the pituitary gland serving as an anatomical reference. Matrix size for T_2_W was 256×256 and 128×128 for the parametric MRI acquisitions.

High-resolution anatomical T_2_W images were obtained with a rapid acquisition relaxation-enhanced (RARE) sequence, with the following parameters: TR = 2500 ms, TE = 26 ms, number of excitations (NEX) = 4.

T_2_ mapping was performed using a multi-slice multi-echo (MSME) sequence with TR = 5000 ms, 50 echoes, TE = 12-600 ms, and NEX = 1.

T_2_* mapping employed a multi-gradient echo (MGE) sequence with TR = 330 ms, 10 echoes, TE = 2-38 ms, flip angle = 30° and NEX = 8.

MTR maps were calculated from MTI sequences. Paired spin echo (SE) images were acquired: one with an MT pulse (MT_on_) and another without (MT_off_), both with TR = 2500 ms, TE = 10 ms and NEX = 1. The MT_on_ sequence incorporated a train of 50 RF pulses (bandwidth = 550 Hz, length = 5 ms, power = 5.5 µT, offset = 1500 Hz).

DTI acquisitions were performed using a Stejskal–Tanner sequence with a four-segment echo-planar (EPI) readout. Diffusion gradients were applied in 15 directions, with 3 b_0_ images, 2 b values of 400 and 1800 s/mm^2^ in each direction, TR = 3000, TE = 32.5ms, Δ = 20 ms, and δ = 4 ms.

### 3.3 Processing details

The studies were processed using Resomapper, applying the Patch2Self denoising filter for DTI studies. All filter parameters and other specifications were kept at their default values as defined by the software. Subsequently, a region-of-interest (ROI)-based analysis was conducted to examine sex differences and regional variations across all parameter maps.

For ROI data extraction, all images were first co-registered using ANTsPy (61), generating a T_2_W template for the entire cohort. Manual ROI delineation was performed on this template, covering key brain regions (Fig 6), including the cortex (CTX), hippocampus (HIP), thalamus (TH), and hypothalamus (HY), using the Allen mouse brain atlas as a reference (62). Pixel data from these ROIs were extracted using a custom macro in ImageJ (63) and were subsequently filtered and analyzed as described in the following section. All scripts and macros employed in this study will be made available at Github upon publication.

**Fig 6.**
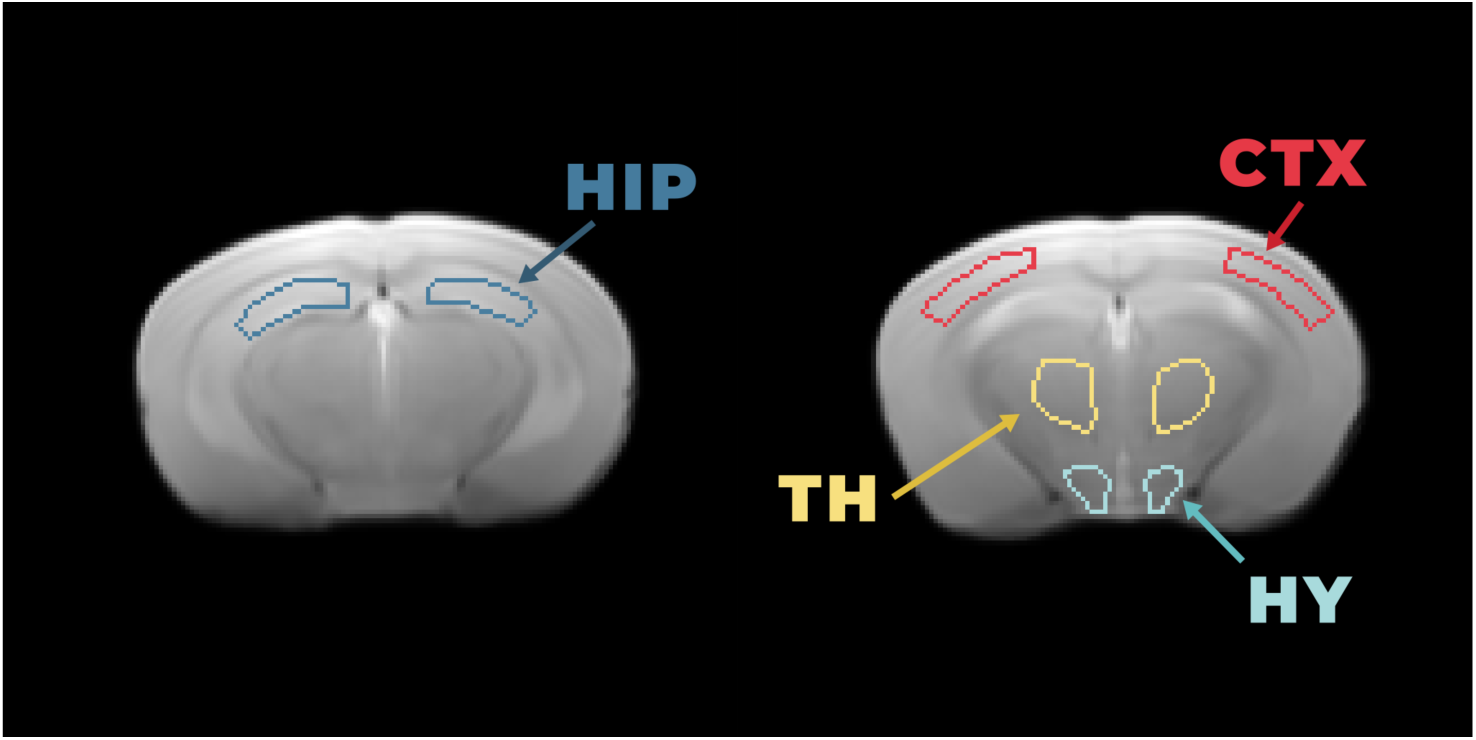
**Example of ROIs selection**. ROIs selected for analysis -cortex (CTX), hippocampus (HIP), thalamus (TH), and hypothalamus (HY)-overlaid on the anatomical T_2_W MRI template averaged across all subjects.

### 3.4 Statistical analysis

The obtained ROI data was analyzed using R software (64). First, the pixel values of each parametric map within each brain region were automatically filtered to exclude extreme outliers, defined as values lying outside the range of the 1st or 3th quartile ± 1.5*interquartile. Subsequently, the mean value for each ROI was computed, and statistical analysis was performed using linear mixed effect (LME) models (65,66).

For each MRI parameter, a model was constructed incorporating the fixed effects of sex, brain region, and their interaction, as well as a random effect accounting for inter-subject variability (subject-level random intercept). Models were developed and fine-tuned based on the Bayesian Information Criterion (BIC) and residual distribution analysis. The statistical significance of fixed effects was assessed within this framework, followed by post-hoc pairwise contrasts where relevant. Multiple comparisons were corrected using the False Discovery Rate (FDR) method.

## 4. Results and discussion

Demonstrating the capabilities of the Resomapper software validates its usefulness in imaging workflows. It enabled the generation of high-quality parametric maps from T_2_, T_2_*, MTR and DTI acquisitions, ensuring accurate pixel-wise model fitting and facilitating quantitative regional analyses. Using the established pipeline also helped in the standardization of the process, saving time in comparison to previous workflows and facilitating the supervision of the results. Fig 7 shows the design of the complete experimental and processing workflow.

**Fig 7.**
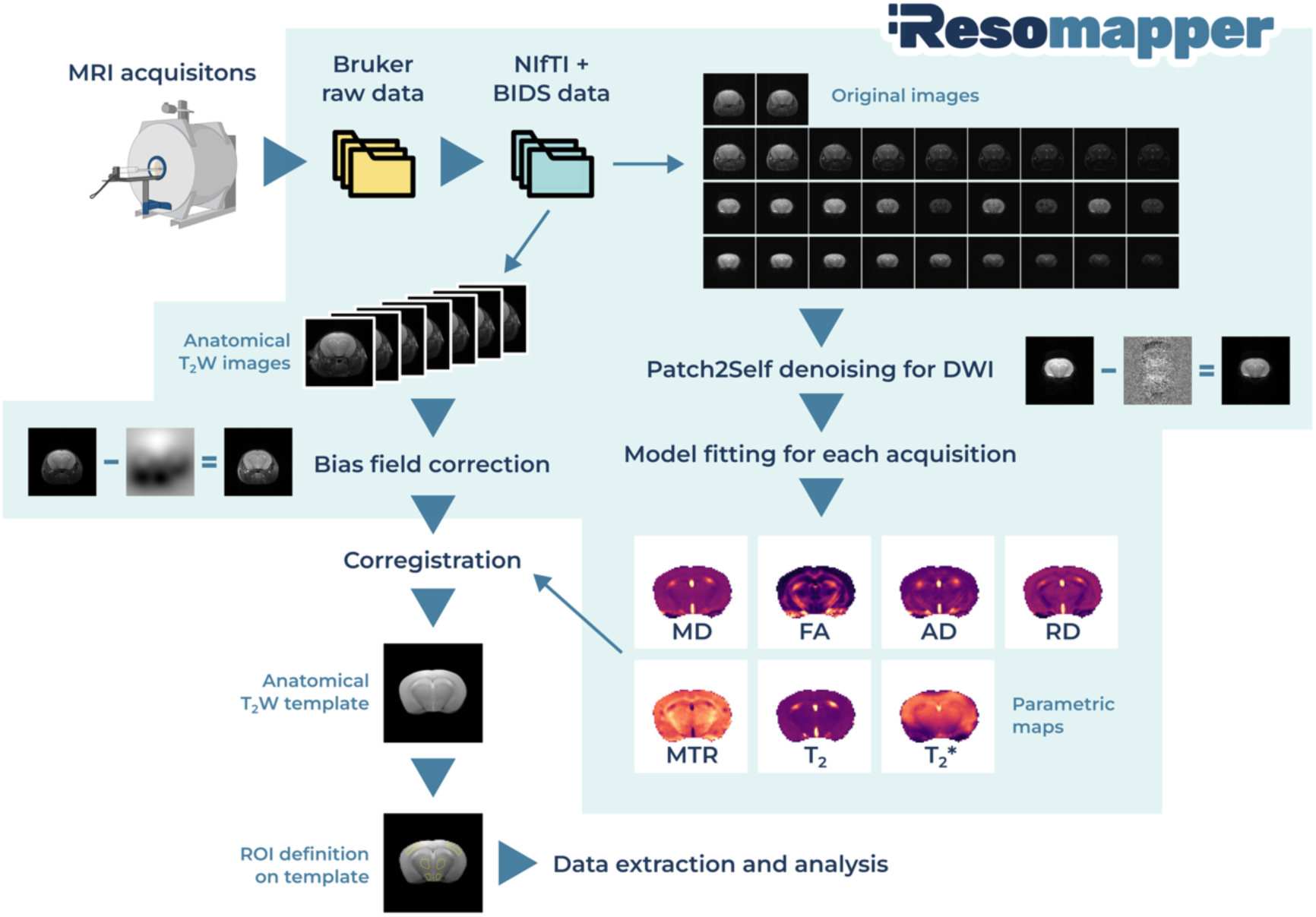
**Processing pipeline using Resomapper**. Representative pipeline applied in this work from the original acquisitions to the final analysed values.

The preclinical example presented in this study illustrates its application in processing multiparametric MRI data, enabling the exploration of potential sex-related differences in MRI parameters, and providing representative values obtained for different brain regions in the mouse, obtained from standard sequences. The mean MRI parametric results for each experimental group and brain areas are depicted on Table 1 and graphically represented on Fig 8 and can be used as a healthy reference for future mice brain studies.

**Table 1.**
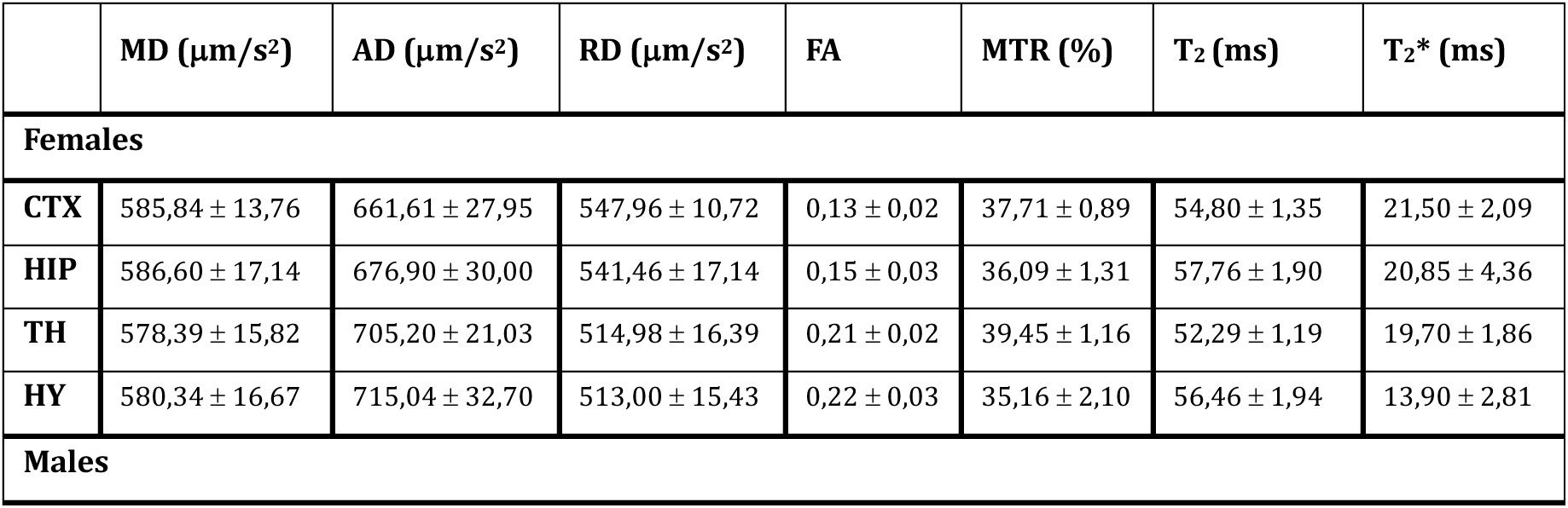
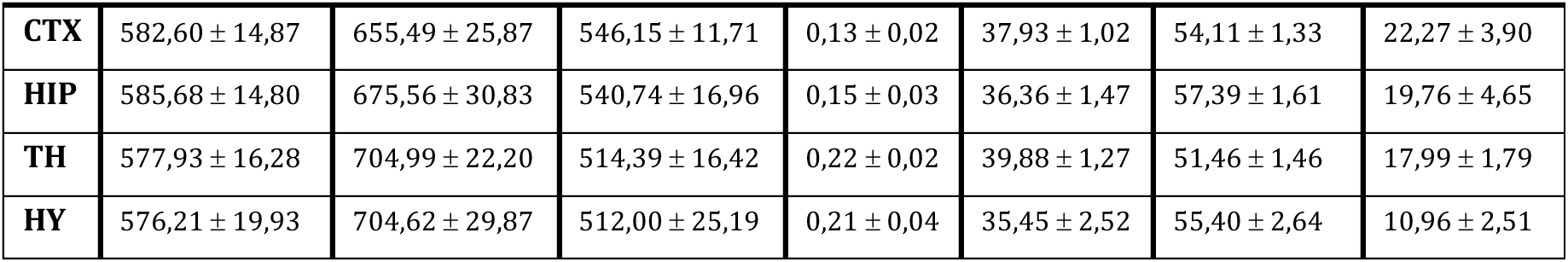
Mean and standard deviation (SD) of the obtained MRI parameters for each brain area of all the subjects of the different sexes.

**Fig 8.**
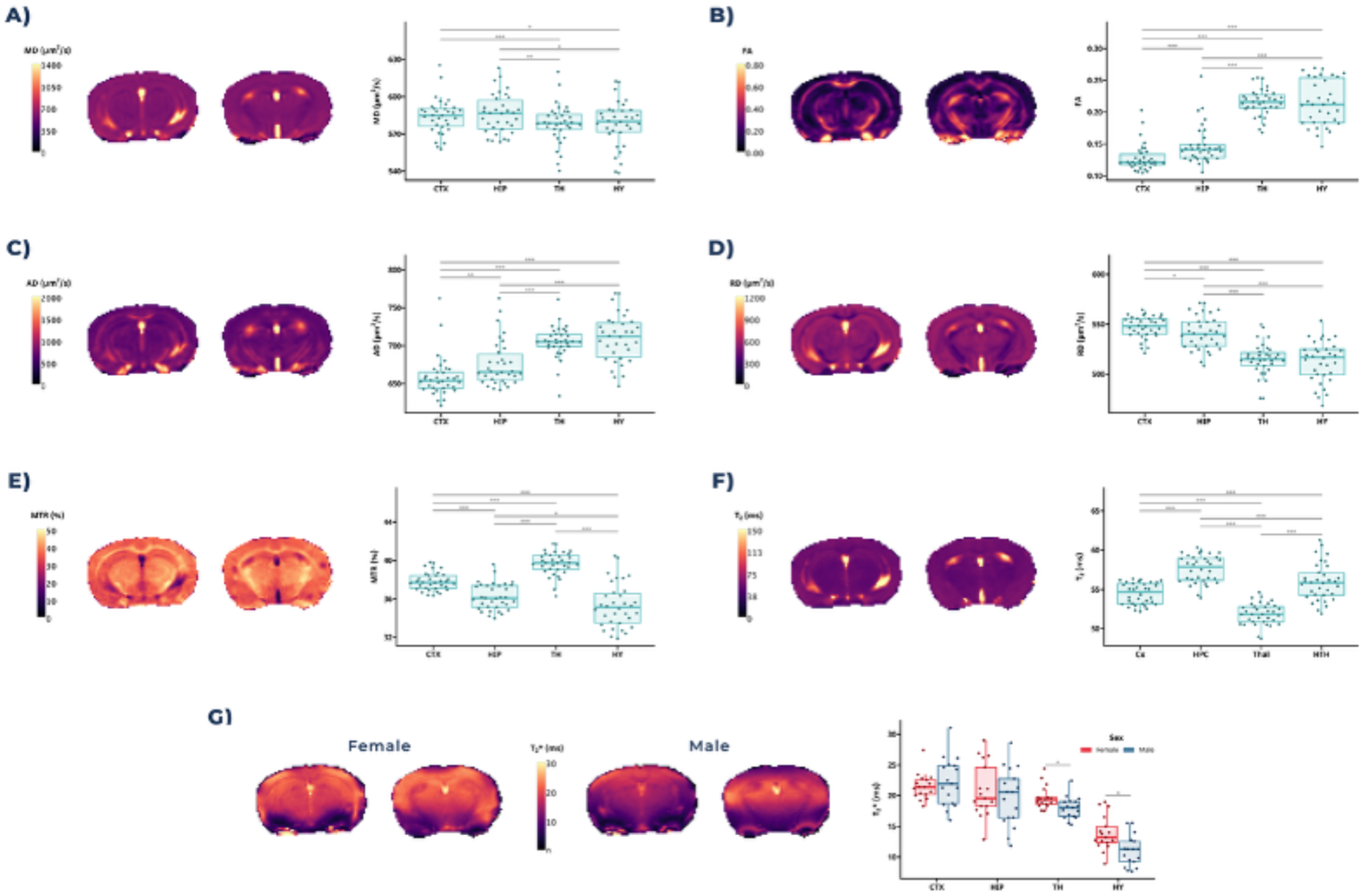
Boxplot graphs of the parametric MRI results by brain area. An illustrative map of one subject is shown next to the graphs, except in G), where one map of a female subject and one of a male subject are shown, as T_2_* results show differences between sexes in the thalamus and hypothalamus.

A significant effect of brain area was observed for all parameters, reflecting inherent differences in tissue microstructure. Regarding DTI parameters, MD exhibited only subtle variations between areas likely due to averaging of diffusivity across all directions, which tends to dampen regional contrasts. In contrast, FA displayed the greatest ability to discriminate between brain areas, with the highest FA values in the TH and HTH, and lowest in the CTX and HIP. AD followed a similar pattern to FA, showing increased values in regions where diffusion movement is more directionally constrained, such as the TH and HTH. Oppositely, RD values were lower in these same areas, in agreement with the predictions of the diffusion tensor model, where higher anisotropy is associated with reduced perpendicular diffusivity.

All studied brain areas correspond to grey matter (as is the case for most of the mouse brain) and, accordingly, display lower FA values compared to white matter. This is expected, as the directionality of diffusion in grey matter is generally less pronounced. Nevertheless, each region possesses distinct organizational and cellular compositions, resulting in the observed DTI differences. The CX is characterized by a highly complex and heterogenous cytoarchitecture, including numerous neuronal bodies, extensive dendritic trees, and local circuitry with less uniform fiber orientation (67). These features together contribute to the disorganized movement of water molecules and hence lower FA. The HIP exhibits a laminate microstructure and a high density of astrocytes and microglia (67,68). Its intrinsic complexity and varying fiber orientations also results in moderately low anisotropy values. In contrast, the TH, and to a lesser extent the HTH, are grey matter structures that, present relatively greater connectivity and contain small but coherent fiber bundles connecting to other brain regions. This underlying microstructural organization leads to higher FA values in these areas relative to the CTX and HIP, despite their classification as grey matter.

In terms of MTR, the TH exhibits the highest values, indicating greater macromolecular content, followed by the CTX. The HY shows the lowest MTR values. T_2_ values display an inverse but complementary pattern to MTR: regions with lower macromolecular content correspondingly have higher T_2_ values, reflecting a higher water content. These findings align with the previously described tissue characteristics: more densely packed areas, such as the CTX and TH, are readily distinguishable, particularly when using MTR measurements (69).Regional differences in T_2_* values reveal a gradient along the vertical axis of the brain, with higher values observed in the upper areas and progressively lower values toward the lower regions, with the HTH exhibiting the lowest T_2_* values. This pattern likely reflects variations in tissue composition and microenvironment, as T_2_* relaxation is sensitive to magnetic susceptibility effects induced by local iron content, myelination, and tissue microstructure. The hypothalamus is known to have relatively higher iron concentration and denser cellular packing compared to other gray matter regions, which can cause increased magnetic field inhomogeneities and consequently shorter T_2_* relaxation times.

Beyond these regional, our analysis uncovered a sex-related effect exclusively to T_2_* values, with female mice showing higher T2* in the TH and HTH compared to males. This finding may potentially be attributed to differential physiological responses to anesthesia between males and females. Isofluorane is known to induce various cerebrovascular effects, including alterations in cerebral blood flow (70) and metabolic suppression (71). Such effects can be detected in T_2_*W images (71–73), due to the sensitivity of T_2_* to magnetic field inhomogeneities arising from deoxyhemoglobin, unlike from oxyhemoglobin which is diamagnetic. The physiological impact of anesthesia is not only dose-dependent but also influenced by sex. Several studies indicate that female subjects exhibit greater resistance to the hypnotic effects of anesthesia, and recover faster than males (74,75). Research in mice has further shown that hypothalamic nuclei involved in sleep, which are sexually dimorphic and hormonally regulated, exhibit more pronounced neuronal activity in anesthetized males, as measured by c-Fos immunofluorescence assays (74). Taken together, these factors complicate the interpretation of our observed T_2_* differences. While a sex-dependent differential effect of anesthesia on hypothalamic activity is a plausible contributing factor, further investigation is necessary to draw definitive conclusions. Additionally, since the T_2_* sequence was the last acquired in the MRI protocol, cumulative anesthetic effects may have been more pronounced at this stage.

In summary, these findings underscore the importance of including both sexes in preclinical and clinical studies and highlight the need for caution when interpreting results that may be influenced by anesthesia-related effects.

## 5. Conclusion

This work presents Resomapper, demonstrating its feasibility for standardized multiparametric qMRI analysis. By integrating various processing tools into a single, user-friendly framework, Resomapper streamlines qMRI workflows and promotes reproducible, high-quality image analysis. Its support for diverse preprocessing techniques, multiple imaging modalities, and standardized data formats makes it a valuable tool for researchers with varying levels of programming expertise.

Future developments will focus on expanding modality support, improving automation, and enhancing compatibility with clinical datasets. Resomapper is actively maintained by the authors, and additional functionalities are planned for implementation in the near future, such as automatic mask definition, the inclusion of more pre-processing options (e.g. motion correction), and the ability to process further modalities like perfusion MRI. The results from the preclinical application example provided in this work can serve as a reference for users intending to apply Resomapper to mouse studies, offering expected values for standard MRI parameters in healthy mice. Furthermore, these results underscore the importance of including both sexes in preclinical research and highlight potential confounding effects of anesthesia in MRI results, particularly in T_2_*, which should be considered when designing and interpreting MRI studies.

## Data Availability Statement

All code used for this article will be available upon publication. The experimental data presented in this study are available on request from the corresponding author due to the need for a formal data sharing agreement.

## Acknowledgements

The authors are indebted to the staff of the Biomedical NMR Facility Sebastián Cerdán for their continuous technical support.

## Author Contributions

Conceptualization: BL and PLL. Data Curation: RGA, BL and PLL. Formal Analysis: RGA, AF and BL. Funding Acquisition: BL and PLL. Investigation: RGA; NAR and PL. Methodology: RGA, AF, NAR, BL and PLL. Project Administration: NAR, BL and PLL. Resources: BL and PLL. Software: RGA. Supervision: NAR, BL and PLL. Validation: RGA, AF, NAR, BL and PLL. Visualization: PLL. Writing - Original Draft Preparation: RGA; Writing - Review & Editing: RGA, AF, NAR, BL and PLL.

All authors have read and agreed to the published version of the manuscript.

The authors declare no competing interests that could be perceived as influencing this work.

## Funding

This work was supported in part by grant PID2021-122528OB-I00 (MICINN/AEI/FEDER, UE) awarded to PLL, PID2021-126888OA-I00 (MICINN/AEI/FEDER, UE) to BL, scholarship PIPF-2022/SAL-GL-25871 to RGA and PRE2022-105662 to AF.

## Institutional Review Board Statement

All experimental procedures were approved by the Ethics Committee of the Community of Madrid (PROEX 288.4/22; approved 18 November 2021) and follow the national (R.D.53/2013) and European Community guidelines (2010/62/UE) for the care and management of experimental animals.

## Notes

### Competing Interest Statement

The authors have declared no competing interest.

